# RPA resolves conflicting activities of accessory proteins during reconstitution of Dmcl-mediated meiotic recombination

**DOI:** 10.1101/356592

**Authors:** Yuen-Ling Chan, Annie Zhang, Benjamin P. Weissman, Douglas K. Bishop

## Abstract

Dmc1 catalyzes homology search and strand exchange during meiotic recombination in budding yeast and many other organisms including humans. Here we reconstitute Dmc1 recombination *in vitro* using six purified proteins including Dmc1 and its accessory proteins RPA, Rad51, Rdh54/Tid1, Mei5-Sae3, and Hop2-Mnd1 to promote D-loop formation between ssDNA and dsDNA substrates. Each accessory protein contributed to Dmc1’s activity, with the combination of all six proteins yielding optimal activity. The ssDNA binding protein RPA plays multiple roles in stimulating Dmc1’s activity including by overcoming inhibitory effects of ssDNA secondary structure on D-loop reactions, and by stabilizing and elongating D-loops. In addition, we demonstrate that RPA limits inhibitory interactions of Hop2-Mnd1 and Rdh54/Tid1 that otherwise occur during assembly of Dmc1-ssDNA nucleoprotein filaments. Finally, we report interactions between the proteins employed in the biochemical reconstitution including a direct interaction between Rad51 and Dmc1 that is enhanced by Mei5-Sae3.

## INTRODUCTION

During meiosis, chromosomes undergo high levels of homologous recombination. Meiotic recombination generates genetic diversity as well as the reciprocal crossover chromosomes required to promote high fidelity reductional chromosome segregation. Meiotic recombination is initiated by the transesterase Spo11 which generates double-stranded DNA breaks (DSBs)(1, 2). DNA ends created by DSBs are processed by nucleases to form 3’ overhanging tracts of ssDNA. The strand exchange protein Dmc1, assembles de-novo on these ssDNA ends, via its high affinity binding site (Site I), with the help of a subset of its accessory proteins. Dmc1 then searches for homologous duplex DNA sequences in a process involving its low affinity binding site (Site II) and carries out strand exchange to form tracts of hybrid dsDNA in which the incoming Dmc1-bound ssDNA and the complementary strand of the original duplex are basepaired. The ‘like” strand of the original duplex is displaced as ssDNA, forming a displacement loop (D-loop).

Replication protein A (RPA) is an essential and abundant protein with no enzymatic activity. It is composed of three subunits: RPA1 (70 kd), RPA2 (30 Kd), and RPA3 (14 kd). RPA has four well defined DNA binding domains that cooperate to bind ssDNA tightly and selectively (relative to dsDNA) with affinity of ~ 10^10^ M^−1^ (3, 4). *In vivo*, binding of RPA to ssDNA is known to play key roles in a wide array of DNA metabolic pathways including DNA replication, nucleotide excision repair, base excision repair, mismatch repair, mitotic repair by homologous recombination, and meiotic recombination. However, the mechanisms through which RPA contributes to these pathways is not fully understood, particularly in the case of meiotic recombination (5-8).

In addition to RPA, genetic and biochemical studies have provided evidence for key accessory proteins that stimulate the activity of Dmc1 (9-18). The main accessory proteins include Rdh54 (a.k.a. Tid1), Rad51, and two heterodimers Mei5-Sae3 and Hop2-Mnd1. All these proteins bind both ssDNA and dsDNA. Rdh54/Tid1 is a dsDNA-specific, ATP hydrolysis-dependent, DNA translocase (19). Rdh54/Tid1and its functionally redundant paralogue Rad54 prevent sequestration of Rad51 and Dmc1 by displacing them from toxic non-recombinogenic complexes formed by direct binding to dsDNA (20-22). Rdh54/Tid1 is also a functionally redundant paralogue of Rad54 which is thought to serve as a “heteroduplex pump” that stabilizes nascent D-loops while displacing the cognate strand exchange protein from strand exchange products (23). Mei5 and Sae3 form a heterodimer that stimulates Dmc1 filament assembly on RPA-coated ssDNA (13, 15). *In vivo*, Mei5-Sae3 is strongly required for DSB-dependent immunostaining foci formed by Dmc1 and for Dmc1-dependent D-loop formation, as assayed by 2-D gels and Southern blotting at a meiotic recombination hotspot (24). Hop2 and Mnd1 also functions as a heterodimer with a significant preference for binding dsDNA over ssDNA. Although Hop2-Mnd1’s *in vivo* function is related to that of Mei5-Sae3 in that it is strongly required for D-loop formation *in vivo*, Hop2-Mnd1 is unique among recombination proteins in that it binds chromosomes to form visible immunostaining foci that are completely independent of DSBs and display low levels of colocalization to the foci formed by Dmc1 (17, 18). Finally, Rad51 is a structural and functional relative of Dmc1 that plays the direct catalytic role in mitotic recombination (25, 26). In addition, and particularly important for this study, Rad51 also serves as an accessory protein to Dmc1 during meiosis (11, 27-29). Rad51’s strand exchange activity is inhibited during meiosis and is not required for normal levels of meiotic recombination in budding yeast (30). Rad51 is required for normal recruitment of Dmc1 to DSBs *in vivo;* the Dmc1 foci formed in *rad51* mutants are faint compared to those in wild type nuclei (31, 32). Rad51 has also been shown to be capable of enhancing Dmc1’s D-loop activity in conjunction with Mei5-Sae3 (11).

When assayed *in vitro*, individual accessory proteins stimulate Dmc1’s D-loop activity (10, 11, 15, 16, 33-36). However, no reconstitution of the activities of all Dmc1 accessory proteins has been reported. Here we describe a 6 protein (10 polypeptide) *in vitro* reconstitution system involving Dmc1 and 5 accessory proteins that stimulate its activity, RPA, Rad51, Mei5-Sae3, Hop2-Mnd1, and Rdh54/Tid1. Our findings show that RPA is particularly important for the high level of Dmc1 activity in the reconstituted system, with three distinct mechanisms contributing to RPA’s overall activity. Finally, we examine protein-protein interactions associated with D-loop activity and present evidence for a novel interaction between Dmc1 and Rad51 that is enhanced by interactions involving Mei5-Sae3.

## MATERIALS AND METHODS

### Protein and DNA preparations

All proteins used in this study are *Saccharomyces cerevisiae* proteins. RPA was expressed from p11d-sctRPA in *E. coli* and purified as described by Binz *et. al.(37).* His6-tagged Dmc1, Mei5-Sae3, Rad51-II3A, Rdh54/Tid1, and Hop2-Mnd1 (16), were expressed and purified as described previously (11, 38). Supercoiled pRS306 and its variant plasmids used for D-loop assays were purified without DNA denaturation using triton lysis buffer and fractionated by cesium chloride density gradient banding (39). The 90C ssDNA is a 90 nt ssDNA and its sequence is homologous to the DNA sequence at position 764 - 853 in pRS306. The 90NF is a 90 nt ssDNA (sequence 5’GG CAC CAACACAAAACACAT CTACACT CAACAAT CACTTTTTAT ACAACACTT CTT CTCTCACATACAACACTTCTGGCACCAACACAAA) and it was transposed from an RNA sequence that was experimentally determined to have no secondary structure above 25 °C (40). The 90NF ssDNA is also predicted to have no secondary structure at 37 °C by the computer programs mFold and OligoAnalyzer tool (IDT). Both 90 nt ssDNA were synthesized by IDT and purified on denaturing gels. The dsDNA version of 90NF was inserted into pRS306 replacing the sequence of 90C at exactly the same location (position 764 - 853 in pRS306); pNRB722 was used as the homologous dsDNA template in D-loop assays with 90NF ssDNA. The 563 nt ssDNA used is identical to the nt’s 306 - 868 of the Crick strand of pRS306. In order to obtain the 563 nt ssDNA, a dsDNA fragment tagged with biotin at the 5’ end of one strand was generated by PCR, then bound to Dynabeads MyOne Streptavidin (Invitrogen) following vendor’s instructions. Subsequently, the strand of 563 nt ssDNA without biotin was released from the bound dsDNA using 0.4 N NaOH with incubation on ice for 2 min. The released 563 nt ssDNA was immediately neutralized upon strand separation and recovered by ethanol precipitation. The ssDNAs were 5’-^32^P labeled by a standard protocol and the unreacted g-^32^P-ATP in the labeling reactions was removed by G25 microspin column (GE Healthcare), and residual T4 polynucleotide kinase was heat-inactivated. Desthiobiotin-90C ssDNA is a variant of the 90C oligo that carries a desthiobiotin moiety with a TEG spacer arm at its 5’ end, and was synthesized and purified by IDT.

### Antibodies

Antibodies against purified Mei5-Sae3, Rad51-II3A, Hop2, and Rdh54/Tid1 were raised in rabbits, antibodies against purified Dmc1 were raised in a goat, at the Pacific Immunology Corp. Antibodies against RPA2 (the 30 Kd subunit of RPA), raised in a rabbit, were a gift from A. Shinohara (Osaka University, Japan). Antibodies were used at 1:1,000 to 1:3,000 dilution in Western blotting. Arp7 anti-goat antibody was purchased from Santa Cruz Biotechnology. The Hop2 antibody cross-reacts with Mnd1. The IgG fractions from antisera were affinity purified by protein A or protein G agarose (GE Healthcare) using standard methods.

### D-loop assay

Reactions (10 μl) were carried out at 37 °C in D-loop buffer (25 mM Tris-HCl (pH 7.5), 1 mM DTT, 5 mM MgCl_2_, 3 mM ATP, 0.25 mM CaCl_2_, and 100 μg/ml BSA).

Concentrations of proteins and their abbreviations are D = Dmc1 (3 μM), A = RPA (0.2 μM), M = Mei5-Sae3 (0.5 μM), R = Rad51-II3A (0.25 μM), H = Hop2-Mnd1 (0.2 μM), T = Rdh54/Tid1 (0.1 μM). ssDNA in the reactions for 90C and 90NF were 30 nM (2.7 μM nt); for 563 nt was 4 nM (2.2 μM nt). dsDNA was supercoiled plasmid pRS306 at 5 nM (22 μM bp) for reactions using 90 mer oligos, or 2.5 nM (11 μM bp) for reactions using the 563nt oligo. Proteins in various combinations were added to reactions using either a pre-mixed or a pre-staged regimen. For the re-mixed regimen, proteins were first mixed on ice before adding ssDNA. After DNA addition, samples were at 37 °C for 5 min to allow formation of nucleoprotein filaments; dsDNA was then added and the reactions incubated for an additional 30 min to form D-loops. For the pre-staged regimen, protein mixtures were added to ssDNA and incubated for 5 min. Hop2-Mnd1 and/or Rdh54/Tid1 were then added as a mixture with dsDNA, followed by a final incubation for 30 min. Hop2-Mnd1 and/or Rdh54/Tid1 were pre-incubated with dsDNA at room temperature for 3 min before adding to nucleoprotein filaments. For both regimens, D-loop formation was terminated by deproteination with the addition of 0.5% SDS and 0.5 mg/ml proteinase K and incubation was for 5 min at 37 °C. 0.2 volumes of loading buffer (25 mM Tris-HCl (pH 7.5), 50% glycerol, 0.1% bromphenol blue, and 0.1% xylene cyanol) was added, and reaction products separated by electrophoresis in 0.9 % agarose gel (10 cm × 14.5 cm) in TAE buffer (40 mM Tris-HCl, pH 7.5, 5 mM sodium acetate, 1 mM EDTA). Electrophoresis was at room temperature at 8 volts/cm for 1 hr 45 min. The gel was dried onto positively charged nylon membranes (Roche), exposed to imaging plate, analyzed using the Typhoon 9200 Imager, and quantified using the computer software Quantity One (BioRad). Both the D-loop band and the unreacted ssDNA band were imaged at the linear range. The D-loop yield was expressed as a percentage of input plasmid DNA. The mean values from three independent trials were plotted for each reaction and error bars show s.e.m.

### Bead “catch and release” assay for DNA binding

RPA and/or Hop2-Mnd1, at concentrations indicated in the legend of Figure 5, were first incubated with desthiobiotin-90C ssDNA (db-ssDNA, 70 nM) in 30 μl D-loop buffer at 37 °C for 8 min to form db-ssDNA-protein complexes (Figure 5A). To capture the db-ssDNA-protein complexes, 2 μl of streptavidin magnetic beads (Roche Diagnostics) was added to the reaction. The bead mixture was incubated for 10 min each at 37 °C and at room temperature on a rotator. The bead bound and unbound fractions were separated using a magnet. The beads were washed two times with 40 μl D-loop buffer containing 0.05% NP40. Protein-ssDNA complexes were then eluted from the beads with 30 μl of biotin buffer (4 mM biotin in 50 mM Tris-HCl, pH 7.5 with 50 mM NaCl) at 37 °C for 13 min. The advantage of the catch and release method compared to standard bead capture assays is that db-ssDNA-protein complexes are eluted from beads to eliminate the fraction of protein bound to beads rather than DNA. Half of the eluted db-ssDNA-protein fraction (15 μΟ was analyzed for protein content by 12% SDS-PAGE and Western blotting. The other half of each eluted fraction was analyzed to determine the yield of eluted DNA by adding an equal volume of urea buffer (8M urea in 50 mM Tris-HCl, pH 7.5, 2 mM EDTA), boiling for 2 min, and running on an 8% urea-PAGE, and staining the gel with SYBR-gold.

### Ni affinity pulldown

Purified Dmc1 (D, 0.5 μM), Mei5-Sae3 (M, 0.5 μM), Rad51-II3A (R, 0.25 μM), Hop2-Mnd1 (H, 0.2 μM), and Rdh54/Tid1 (T, 0.2 μM) were (His)_6_-tagged. Each of these proteins was first incubated individually with untagged RPA (A, 0.2 μM) in 20 μl D-loop buffer at 37 °C for 10 min, followed by the addition of 1.7 μl Ni sepharose beads (30% slurry of Ni high performance sepharose, GE Healthcare). The mixture was then incubated at 4 °C for 30 min on a rotator to capture the interacting protein complexes through the (His)_6_ tag. Beads were pelleted by centrifugation for 0.5 min at 3,000 rpm and washed one time with 100 μl of wash buffer A (25 mM Tris-HCl (pH 7.5), 1 mM DTT, 1 mM MgCl_2_, 1 mM ATP, 50 μΜ CaCl_2_, 0.5 M NaCl, 50 mM imidazole, 100 μg/ml BSA, and 0.05% NP40), and three times with 100 μl wash buffer B (wash buffer A with 0.1 M NaCl). The Ni captured proteins were eluted by boiling in 20 μl 1.7%SDS and 0.1 M DTT solution, separated on 12% SDS-PAGE, and analyzed by Western blotting.

### Co-immunoprecipitation

Physical interaction between RPA (0.2 μM) and Dmc1 (0.5 μM), RPA and Mei5-Sae3 (0.5 μM), or RPA and Rad51-II3A (0.25 μM) was tested by co-immunoprecipitation assay. Proteins were mixed (as indicated in Figure 6B, 6C) in a 20 μl reaction containing D-loop buffer and incubated at 37 °C for 10 min. Purified antibodies (0.5 μl against either the RPA2 or Dmc1 were added as indicated and the mixtures were incubated at 4 °C for 1 h on a rotator. Equal volumes of 0.5 μl magnetic-conjugated protein G (Dynabeads protein G, Invitrogen) and magnetic-conjugated protein A (NEB) were added to the mixture, and further incubated at 4 °C for 1 h. The immunoprecipited complexes immobilized on magnetic beads were then washed with 100 μl of wash buffer (25 mM Tris-HCl (pH 7.5), 1 mM DTT, 1 mM MgCl_2_, 1 mM ATP, 50 μM CaCh, and 100 μg/ml BSA, 0.1 M NaCl, and 0.05% NP40) four times. The bead-bound protein was eluted by boiling in 40 μl 1.7% SDS and 0.1 M DTT denaturing buffer, and a 20 μl eluate was loaded onto 12% SDS-PAGE for analysis. Proteins were detected by Western blot using antibodies against all proteins in the reactions. Three independent trials were done for each reaction.

### Quantitative Western analysis of protein levels *in vivo*

200 ml cultures of SK-1 strain DKB1772 (ho::hisG/”, HIS4::LEU2-(NBam)/his4X::LEU2-(NBam)-URA3, leu2::hisG/”, ura3 (DPst-Sma)/”) were prepared for liquid sporulation and sporulation/meiosis was induced using a previously published method (31). Yeast whole cell protein extracts prepared by the TCA method from three independent meiotic cultures by taking culture aliquots at 0, 2, 3, 4, 5, and 7 hours following induction of meiosis. Antibodies used were as described above. Because steady state levels of Arp7 do not change during meiosis, it was used as an internal loading control to normalize band intensity across all time points. A dilution series of purified target protein of known amount was used to construct a standard curve, and the amount of yeast proteins present at different meiotic time points calculated by linear regression. For calculations, we used the median budding yeast diploid cell volume of 86 × 10^−15^ L, and nuclear volume at 7% of the cell volume (41, 42). For the heterodimers, only one protein concentration in the pair was determined. For the values of RPA in Table 1, it is assumed that all proteins are completely localized to the nucleus.

**Table 1.** Concentration of budding yeast proteins during meiosis

| Protein | Meiosis<br>Peak<br>hr | Molecules<br>$\times 10^4$<br>Per cell | Concentration<br>Per cell<br>($\mu\text{M}$) | Concentration<br>Per nucleus<br>($\mu\text{M}$) | Concentration<br>in<br>reconstitution<br>($\mu\text{M}$) | *In vivo*<br>Nuclear<br>Ratio<br>relative to<br>Rdh54/Tid1 | *In vitro*<br>Ratio<br>relative to<br>Rdh54/Tid1 |
| --- | --- | --- | --- | --- | --- | --- | --- |
| Rdh54/Tid1 | 5 | $0.92 \pm 0.4$ | 0.18 | 2.6 | 0.1 | 1 | 1 |
| Rad51 | 4 | $1.8 \pm 0.9$ | 0.36 | 5.1 | 0.25 | 2 | 2.5 |
| Mei5 | 4 | $5.5 \pm 2.2$ | 1.08 | 15.4 | 0.5 | 6 | 5 |
| Dmc1 | 4 | $8.9 \pm 1.4$ | 1.74 | 24.9 | 3 | 10 | 30 |
| RPA | 4 | $39.2 \pm 3.0$ | 7.70 | 110.0 | 0.2 | 42/3* | 2 |
| Hop2 | 4 | $24.7 \pm 2.0$ | 4.84 | 69.1 | 0.2 | 27 | 2 |

\* The ratio of RPA over Rdh54/Tid1 is 3 if RPA is assumed to be uniformly distributed throughout the cell rather than localized to the nucleus (see text).

## RESULTS

### *In vitro* reconstitution of Dmc1 dependent recombination with five accessory proteins

In order to reconstitute Dmc1 mediated recombination *in vitro*, we developed a system in which the influence of 5 accessory proteins on Dmc1’s activity was determined. These proteins included RPA (abbreviated “A” in all figures), Rad51-II3A (“R”), Rdh54/Tid1 (“T”), and two heterodimers Mei5-Sae3 (“M”) and Hop2-Mnd1 (“H”). Rad51-II3A was used in place of the wild type Rad51 protein to ensure that all D-loop activity observed under the various reaction conditions was contributed by Dmc1 and not Rad51. Rad51-II3A retains DNA binding but not strand exchange activity (11). Rad51 and Rad51-II3A stimulate Dmc1’s D-loop activity in cooperation with Mei5-Sae3. The final concentrations of proteins in our reconstitution experiments were arrived at by titration by varying the concentration of one protein, but keeping all other five proteins constant (Supplementary Figure S1). The protein concentration range explored for each protein in the titrations was informed by results of previous studies (11, 16, 34). The concentration titration range for RPA and Dmc1 was above its theoretical saturation value on ssDNA which was estimated using RPA’s binding site size of 25 nucleotides and Dmc1’s binding site size of 3 nucleotides per protomer. The optimal concentrations of proteins used in the experiments described below are listed in MATERIALS AND METHODS and in Table 1.

After determining the optimum concentration at which each of the five accessory proteins stimulated Dmc1, we carried out D-loop reactions in which different combinations of the proteins were mixed on ice prior to addition of the ssDNA substrate (Figure 1A). D-loop reactions were carried out using a variant of the standard two step method in which ssDNA is added to the strand exchange protein, incubated to allow nucleoprotein filament formation in Step 1, and the supercoiled dsDNA substrate then added to initiate D-loop formation in Step 2. In this case, Dmc1 was added to the ssDNA either alone or as a mixture with accessory proteins. Proteins were premixed rather than added sequentially to ssDNA, as was done in our previous studies (11, 16). This regimen was used to better mimic *in vivo* conditions in which formation of ssDNA initiates assembly of recombination complexes (see legend of Figure 1). Using the “premixed” version of the 2-step protocol, we determined D-loop formation using a 90-mer ssDNA oligonucleotide (designated 90C) as ssDNA substrate and a 4.4 kb supercoiled plasmid carrying a sequence identical to 90C (Figure 1B). During the initial analysis of the data it became clear that RPA had a particularly strong stimulatory effect. We therefore present the data from these experiments as a comparison between a set of mixtures that do not contain RPA to a corresponding set of mixtures that do. We found that (1) D-loop formation is completely Dmc1-dependent regardless of whether or not RPA was present (lanes 10 - 16), confirming that none of the other proteins, including Rad51-II3A, contribute significant D-loop activity under any of the conditions analyzed. (2) In the absence of RPA, the yield of D-loops was low (<0.9%) even in the presence of all the other accessory proteins (Left panel, lanes 2 - 9). In the set of reactions that lack RPA, there were no additive effects of combining accessory proteins on the final yield of products, suggesting that the accessory proteins were conflicting with one another. (3) When RPA was included in the same set of reactions, D-loop yields were dramatically higher (range from 2 - 12%) (Right panel, lanes 2 - 9) than those in corresponding reactions lacking RPA. Unlike the reactions lacking RPA, combining accessory proteins increased the yield of D-loops above the level seen with individual accessory proteins and the reaction containing all 6 proteins yielded more D-loop product than any of the partial reactions (Right panel, lane 9). This result indicated that we have identified conditions under which each protein component in the mixture is able to contribute significant stimulatory activity to the total D-loop yield, and this includes individual contributions from Rad51-II3A and Mei5-Sae3 (Supplementary Figure S2).

**Figure 1.**
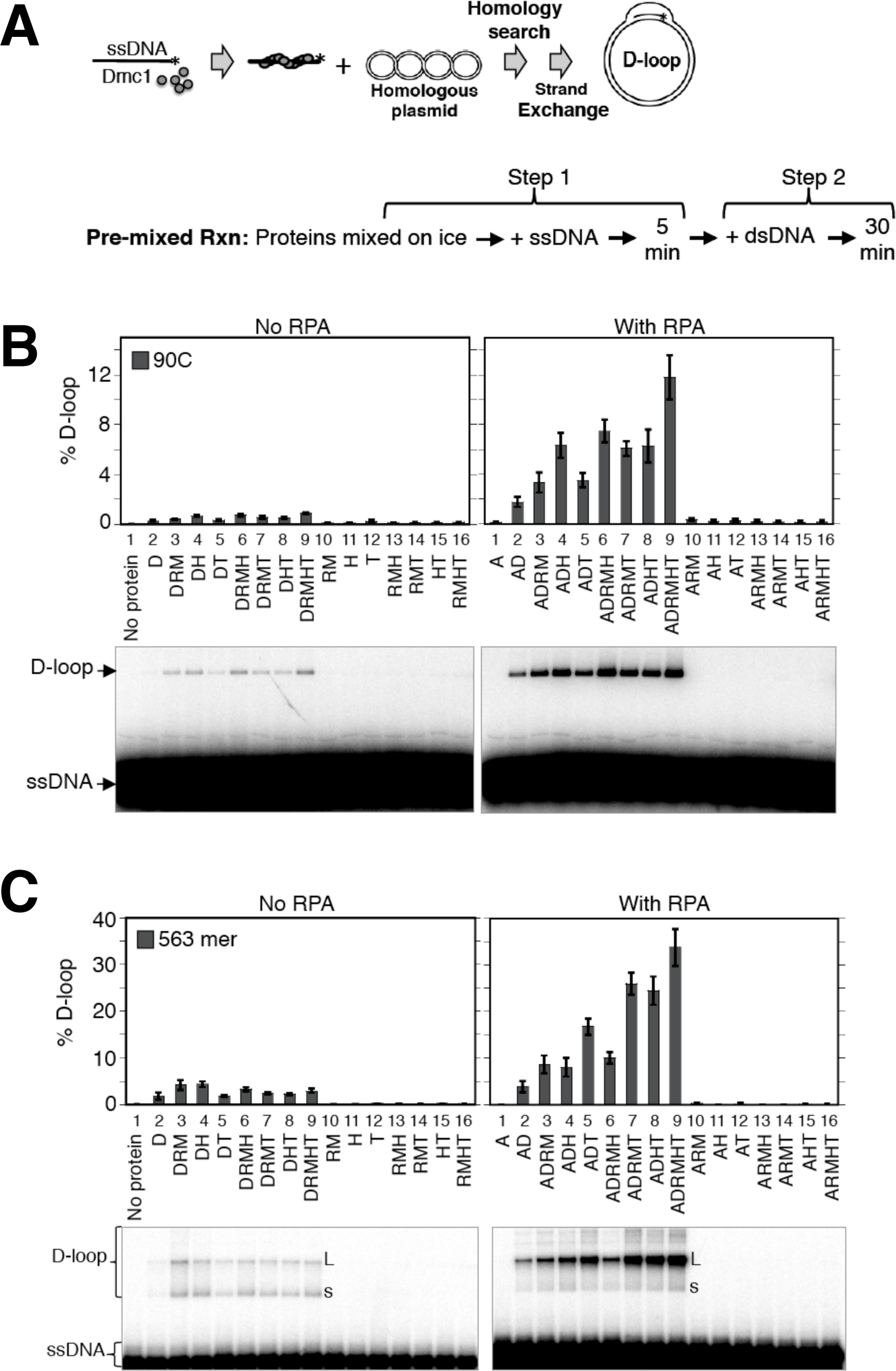
*In vitro* reconstitution of Dmc1-dependent recombination with five accessory proteins. (**A**) Scheme for the 2-step D-loop assay with the pre-mixed reaction regimen. For the first step, reaction components, including pre-mixed proteins, were incubated with ssDNA at 37 °C for 5 min to make Dmc1 filaments. The second step is initiated by addition of target dsDNA followed by incubation for 30 min to form D-loops. (**B**) D-loop assay reactions using the 90C ssDNA (30 nM or 2.7 μM nt) and various mixtures of proteins. Abbreviations for proteins and their concentrations in the reactions: D = Dmc1 (3 μM), A = RPA (0.2 μM), M = Mei5-Sae3 (0.5 μM), R = Rad51-II3A (0.25 μM), H = Hop2-Mnd1 (0.2 μM), T = Rdh54/Tid1 (0.1 μM); dsDNA was supercoiled plasmid pRS306 (5 nM or 22 μM bp). (C) D-loop assay reactions using the 563 nt ssDNA (4 nM or 2.2 μM nt) and various combinations of accessory proteins at the same concentrations as in **B**., dsDNA was at 2.5 nM (11 μM bp). “L” stands for position of the largest D-loops detected, and “s” for position of the smallest D-loops. The D-loop yield was expressed as a percentage of input plasmid DNA. The gel images from D-loop assays were shown and data were quantified and graphed (n = 3, ± s.e.m). [NB: Pre-mixing results in lower levels of D-loop activity than observed when proteins are added sequentially to ssDNA, accounting for the different in D-loop yields seen here as compared to our previously published papers (11, 16)].

To determine if the results obtained with the 90-mer ssDNA substrate would hold for a longer substrate, one similar in length to that of ssDNA tracts that form in the cell (43), we repeated analysis of the activity of protein mixtures using a 563 nt ssDNA (Figure 1C). The results we obtained with the 563-mer were similar to those obtained with the 90C. RPA greatly stimulated D-loop yields regardless of the combination of other accessory proteins present; additive effects of Rad51+Mei5-Sae3, Hop2-Mnd1, or Rdh54/Tid1 were only observed in reactions containing RPA (compare Figure 1C to Figure 1B). As for 90C, the 563-mer substrate yielded the most D-loops when all proteins were present in the reaction with a yield of about 34% (Figure 1C, right panel, lane 9). The relatively high yield of D-loops observed with the 563-mer as compared to the 90-mer was largely dependent on Rdh54/Tid1; reactions containing Rdh54/Tid1 yielded 3 to 4-fold more D-loops with the longer substrate as compared to the shorter one, for which addition of Rdh54/Tid1 only increased yields up to 2-fold (compare Figure 1C to Figure 1B, right panels, lanes 5 to 2, 7 to 3, 8 to 4, and 9 to 6).

A notable observation with the 563-mer was that when RPA was present, D-loop size increased dramatically, as evidenced from the uniformly slow migrating D-loop species observed, as compared to the reactions without RPA, which produced a wider range of D-loop sizes (Figure 1C, lanes 2 - 9, compare both panels). Longer D-loops migrate more slowly on gels because the negative supercoils of the plasmid dsDNA substrate are progressively unwound as D-loops become longer, resulting in slower migration in gels (23). The increase in D-loop size promoted by RPA likely reflects extensive binding of RPA to the displaced strand of nascent D-loops. Such binding is expected to help stabilize nascent D-loops and drive the extension of heteroduplex DNA within those loops (23, 44, 45).

To further characterize the mechanism of stimulation of Dmc1 by the accessory proteins, we carried out kinetic analysis on the 563-mer ssDNA substrate to obtain apparent endpoints and rates of approach to those endpoints for D-loops formation in the presence of Dmc1 and RPA. We find that Rad51+Mei5Sae3, Rdh54-Tid1 and Hop2-, Mnd1 increase both the rate and extent of reaction (Supplementary Figure S3). The results also show that all of the reactions examined are nearing their endpoints by 30 min, the time used to report yields in all other D-loop experiments reported in this paper (Supplementary Figure S4). The results of this kinetic analysis are consistent with the possibility that addition of each protein increases the fraction of ssDNA-protein complexes that is capable of forming stable D-loops. However, the results do not exclude alternative, or more complex, kinetic models.

### Removal of secondary structures from ssDNA by RPA enhances the D-loop forming activity of Dmc1

To determine if the overall stimulation of reactions by RPA involved previously described mechanisms through which RPA, or its prokaryotic functional homolog SSB, stimulates strand exchange reactions. One well known stimulatory activity of RPA, and bacterial SSB, is to remove the secondary structure from ssDNA substrates that otherwise limits the activity of strand exchange proteins (46, 47). We therefore asked if secondary structure was limiting D-loop yield in the absence of RPA, and if the stimulatory activity of RPA could be fully accounted for as an effect of eliminating secondary structure. To answer these questions, we compared the results obtained using the 90C oligonucleotide, which is predicted to have substantial secondary structure at 37 °C, to the results obtained using another 90-mer, designated 90NF (for “not folded”), which is predicted to lack secondary structure under the same conditions (Figure 2). We found that, in the absence of RPA, the D-loop yield was 6-fold higher for 90NF as compared to 90C when all proteins except RPA were added to reactions (Figure 2B, lane 3). These findings suggest that secondary structure of 90C limits the yield of D-loops in the absence of RPA. Importantly, although 90NF is less dependent on RPA than 90C for generation of D-loops; addition of RPA enhances the level of D-loops of 90NF by 2.5-fold from 6 to 15% (Figure 2B, compare lanes 3 and 4), indicating that RPA’s stimulatory activity is not limited to overcoming inhibitory effects of secondary structure. These results indicate that some, but not all, of RPA’s stimulatory activity on 90C results from eliminating the inhibitory effects of secondary structure.

**Figure 2.**
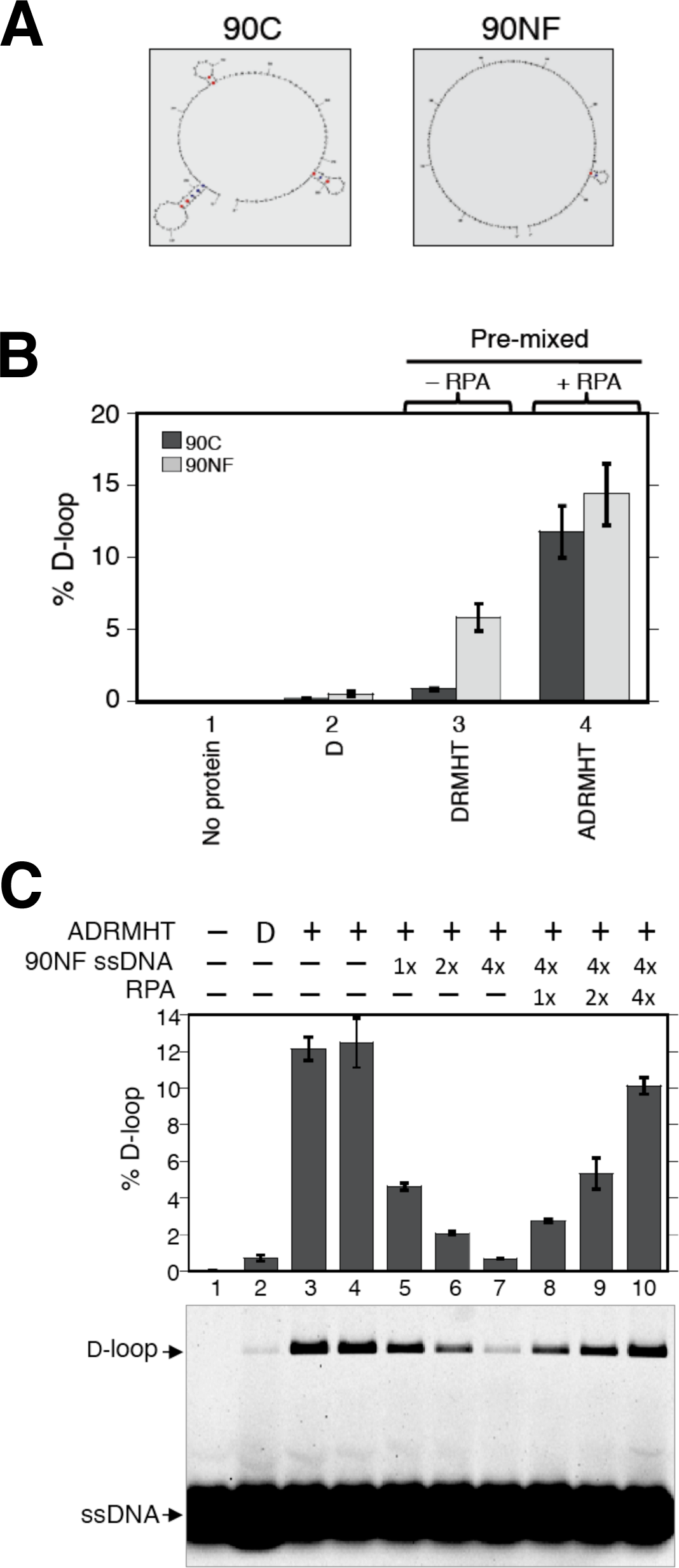
Effects of RPA on secondary structure and inhibitory binding of heterologous DNA to Dmc1-site II. (**A**) Cartoons of predicted secondary structures of 90C ssDNA (significant secondary structure) and 90NF ssDNA (no secondary structure at 37 °C). [NB: the cartoon for 90NF was produced by a computer program using 25 °C rather then 37^°^C as the input temperature because the program does not produce an output image if no secondary structure is detected. The small stem loop shown in the cartoon is not predicted to form at 37 °C.] (**B**) D-loop formation by Dmc1 was examined using 90C ssDNA (30 nM or 2.7 μM nt) and 90NF ssDNA (30 nM or 2.7 μM nt). The pre-mixed regimen was used. Abbreviations and concentrations were as in Figure 1. (**C**) Competition of RPA with Dmc1-site II for ssDNA binding. In the Pre-mixed regimen, all 6 proteins (ADRMHT) were first added to 90C ssDNA (1x = 30 nM or 2.7 μM nt) to make Dmc1-ssDNA filaments. After filament formation, increasing amounts of nonhomologous ssDNA (90NF, 1x = 30 nM or 2.7 μM nt) were added to reactions 5 - 10 as indicated to allow binding to the secondary Site II in Dmc1-ssDNA filaments. RPA (1x = 0.2 μM) was then added in increasing amounts to reactions 8 - 10 before the addition of plasmid pRS306 to each sample to initiate D-loop formation (n = 3, ± s.e.m).

### RPA can outcompete Dmc1’s secondary DNA binding site for ssDNA

In addition to stimulating recombination reactions by removing secondary ssDNA structures, RPA was previously implicated in enhancing strand exchange reactions by binding to the displaced strand of D-loops thereby stabilizing them (44, 45). Although it is technically difficult to directly demonstrate that RPA binds the displaced strand, an indirect approach was used previously to infer this activity. The secondary low affinity DNA binding site (Site II) of *E. coli* RecA is known to bind the displaced (a.k.a. “outgoing”) ssDNA strand briefly before it is released from the interior of the filament following strand exchange. The indirect approach uses a non-homologous ssDNA to inhibit strand exchange by binding to Site II. Subsequent addition of single strand binding protein to the inhibited reaction competes for the heterologous ssDNA and restores D-loop activity (45). In our reconstitution system, 90C ssDNA produced ~ 12% D-loops at 0.2 μM RPA (Figure 2C, lanes 3 and 4). Addition of a 4-fold excess of a non-homologous 90NF ssDNA almost completely inhibited D-loop formation (lane 7). This inhibition was relieved when additional RPA was added to reaction mixtures (lanes 8 - 10). These results indicate that RPA can outcompete Site II for ssDNA. The results of this competition experiment are consistent with our finding that RPA increases D-loop size for the 563-mer substrate and provide additional evidence that RPA enhances Dmc1 D-loop yield by binding to the displaced ssDNA strands of nascent D-loops.

### Reconstituted Dmc1 reactions lacking RPA are more efficient when Hop2-Mnd1 and Rdh54/Tid1 are added late to reactions

Although the results presented above provide evidence for two mechanisms through which RPA enhances Dmc1’s D-loop activity in reconstituted reactions, neither of these mechanisms explains how RPA enables the cooperation of accessory proteins seen in the pre-mixed protocol described above. Previous *in vivo* studies suggest that the Dmc1 stimulatory activities of Rdh54/Tid1 and Hop2-Mnd1 are likely to involve a mechanism acting after the assembly of Dmc1 filaments; Dmc1 focus formation does not depend on Hop2-Mnd1 or Rdh54/Tid1 *in vivo.* Furthermore, previous studies showed that Hop2-Mnd1’s stimulation of Dmc1 activity is optimal only when the protein is added to reactions late, along with the dsDNA template (Figure 3A)(10, 16, 48), suggesting that its presence during Dmc1 filament assembly may have an inhibitory effect. To determine if Rdh54/Tid1 displays similar sensitivity to order of addition, we carried out a set of reactions in which Rdh54/Tid1 was added at different stages (Figure 3B). As for Hop2-Mnd1, Rdh54/Tid1 stimulatory activity was 16-fold greater if the protein was added at the second step of the 2-step D-loop protocol, along with the dsDNA target plasmid, as compared to when it was added at the first step, in a mixture with Dmc1, to ssDNA (Figure 3B; compare reactions 1 and 2). The relatively low yields of D-loops formed when Hop2-Mnd1 or Rdh54/Tid1 was added at the first step as compared to being added at the second step are likely to reflect inhibitory interactions of Hop2-Mnd1 or Rdh54/Tid1 with ssDNA, and/or binding to secondary structures formed by folding of ssDNA. If this were the case, such an inhibitory interaction might limit the ability of Hop2-Mnd1, or Rdh54/Tid1, to cooperate with each other, or with the other accessory factors, for stimulation of Dmc1’s activity. To test this, we delayed the time of addition of Hop2-Mnd1, or Rdh54/Tid1, to protein mixtures containing Dmc1 and Rad51+Mei5-Sae3, but lacking RPA. Using this “pre-staged” regimen, we found that delayed addition of Hop2-Mnd1 or Rdh54/Tid1 increased the yield of D-loops 4.4 fold from about 1.4% to about 6% (Figure 3C). This result indicates that, in the absence of RPA, early addition of Hop2-Mnd1 or Tid1 results in less efficient stimulation than occurs when proteins are added with the dsDNA substrate at the second step of the two-step D-loop protocol.

**Figure 3.**
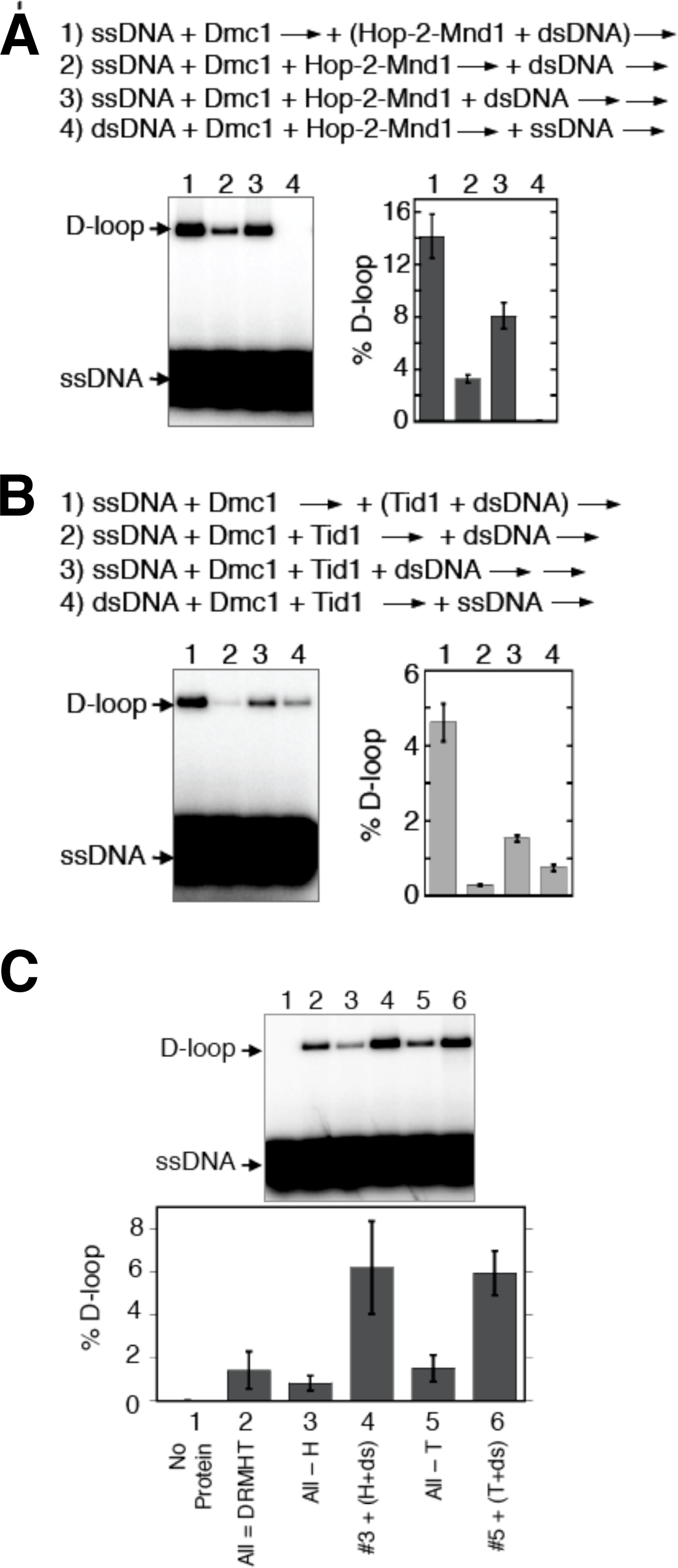
Dmc1 reactions lacking RPA are more efficient when Hop2-Mnd1 and Rdh54/Tid1 are added late to reactions. (**A**) The order of addition of reaction components is as indicated. The concentrations of reactants are as detailed in MATERIALS AND METHODS. The first arrow indicates incubation was at 37 °C for 10 min followed by the second arrow, which indicates an additional incubation at 37 °C for 15 min. Hop2-Mnd1 and dsDNA in reaction 1 were incubated at room temperature for 3 min prior to adding to Dmc1-ssDNA filaments. The gel images from D-loop assays are shown and results graphed (n = 3, ± s.e.m). Data shown here was published previously (16) and are presented for comparison to Figure 3B with permission [NB: permission pending]. (**B**) The same experimental procedure as in **A** except Rdh54/Tid1 was used in place of Hop2-Mnd1. (**C**) Deletion and add-back reactions in the Pre-mixed regimen of Hop2-Mnd1 or Rdh54/Tid1 in the absence of RPA. The order of addition of components were as indicated. Abbreviations for proteins and protein and DNA concentration are as in Figure 1.

### Adding Hop2-Mnd1 and Rdh54/Tid1 at the second step of the D-loop assay reduces the impact of RPA on D-loop yield

If pre-staging the order of protein addition in reconstitution reactions avoids inhibitory interactions between Hop2-Mnd1 and ssDNA and/or between Rdh54/Tid1 and ssDNA, during Dmc1 filament formation, and if RPA reduces or eliminates such inhibitory interactions in the pre-mixed protocol, then RPA’s stimulatory activity is predicted to be less pronounced in the pre-staged protocol as compared to the pre-mixed protocol. To test this prediction, we compared the relative impact of adding RPA to pre-mixed vs. pre-staged D-loop reactions using both 90C and 90NF. As described above (Figure 2B), RPA improved yields in the pre-mix protocol by 13-fold stimulation for 90C and 2.5-fold for 90NF. The amount of stimulation by RPA in the pre-staged reaction was less dramatic than that in the pre-mixed reaction; addition of RPA increased D-loop yield 2-fold for 90C and about 1.3-fold for 90NF (Figure 4, lanes 3 and 4). The results support the conclusion that a substantial amount of RPA’s stimulatory activity in the pre-mixed regimen involves eliminating inhibitory interactions of Hop2-Mnd1 and Rdh54/Tid1 during pre-synaptic filament formation. On the other hand, our results clearly show that addition of RPA increased the yield of D-loops in the pre-staged regimen even for 90NF; therefore, reducing inhibitory interactions during Step 1 and eliminating inhibitory effects of secondary structure, does not fully account for the stimulatory activity of RPA. The residual stimulatory activity seen in the pre-staged reactions for 90NF is likely accounted for by binding of RPA to the displaced strand of nascent D-loops as described above.

**Figure 4.**
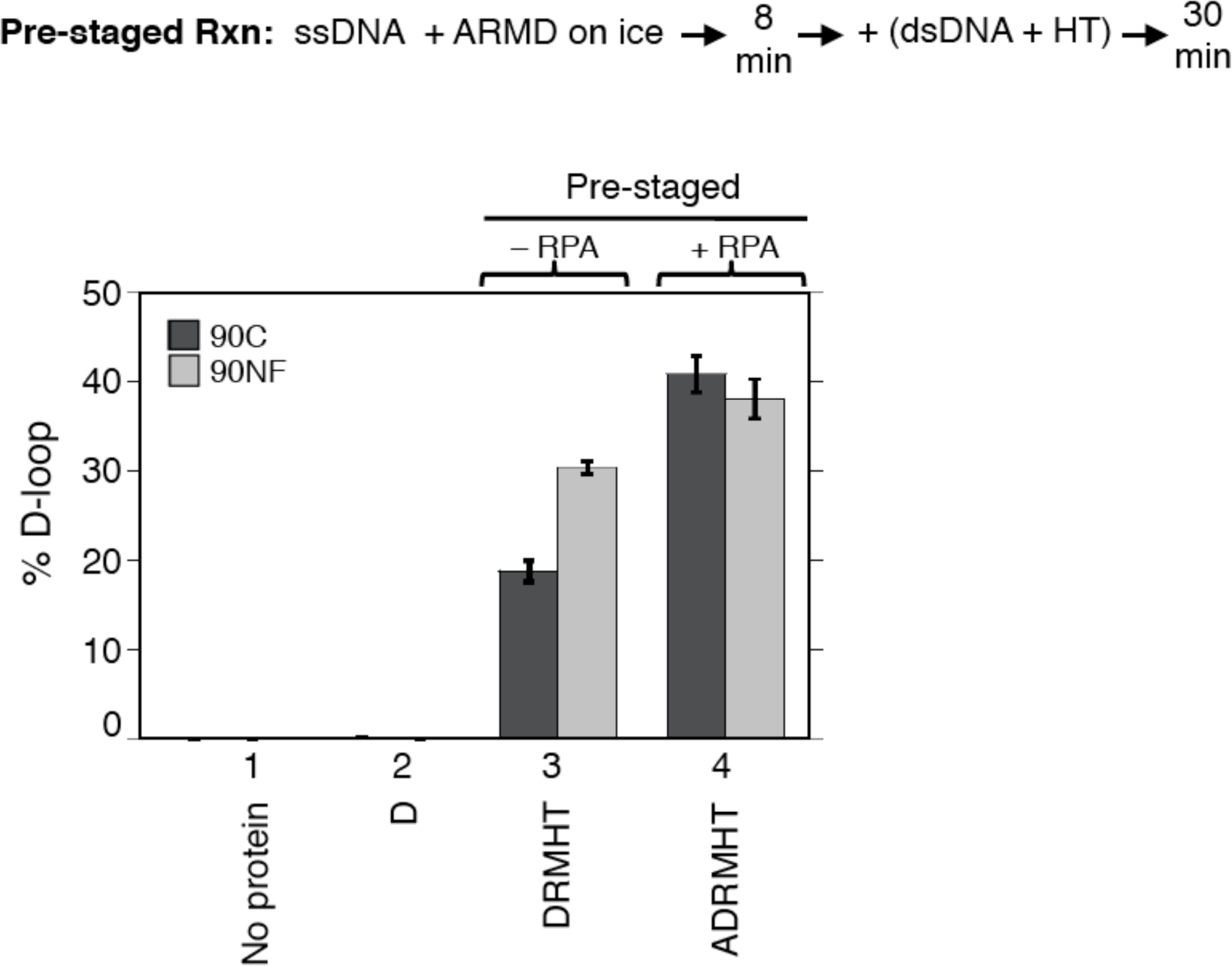
Effect of RPA in pre-staged reactions. D-loop formation by Dmc1 and its accessory proteins was examined using 90C ssDNA and 90NF ssDNA (30 nM or 2.7 μM nt). Reactions were with or without RPA as indicated. Abbreviations and reaction component concentrations were as in Figure 1. (n = 3, ± s.e.m).

### RPA outcompetes Hop2-Mnd1 binding to ssDNA

The finding that early addition of Hop2-Mnd1, or Rdh54/Tid1, limits the yield of D-loops in the absence of RPA raised the obvious possibility that the mechanism through which RPA promotes cooperation of Hop2-Mnd1 and Rdh54/Tid1 involves competition of RPA with the accessory proteins for binding to ssDNA. RPA is well known to bind ssDNA with sub-nanomolar affinity, while Hop2-Mnd1 binds less tightly (49, 50). To confirm this, we used a bead “catch and release” method to measure protein binding to DNA (Figure 5A). This method allowed us to determine: (1) if Hop2-Mnd1 binds the ssDNA substrate under our reaction conditions and (2) if addition of RPA blocks Hop2-Mnd1 binding to ssDNA under the same conditions. Proteins were allowed to bind a variant of the 90C ssDNA oligo that carries a desthiobiotin moiety at its 5’ end (designated as db-ssDNA). Following binding reactions, protein bound oligos were mixed with streptavidin magnetic beads. Unbound protein and oligo were then removed by washing the beads with buffer, and biotin added to release DNA from beads, along with bound protein. The amount of DNA-bound protein was determined by Western blotting, and the amount of DNA by urea-PAGE. As shown in Figure 5B, Hop2-Mnd1 bound ssDNA when added alone, and addition of RPA to binding reactions blocked Hop2-Mnd1 binding. Analysis of unbound proteins (Figure 5C) confirmed this conclusion. No protein was detected in a no DNA control (lane 13), highlighting the advantage of our catch and release method; it avoids background caused by non-specific binding of proteins to beads. These results support our model that RPA enhances reconstituted reactions by blocking inhibitory interaction of Hop2-Mnd1 with ssDNA. We were unable to carry out equivalent catch and release experiments for Rdh54/Tid1 presumably because binding of Rdh54/Tid1 was too weak to withstand the washing steps required for the method, but we were able to demonstrate by fluorescence polarization assay that the apparent binding affinity of Rdh54/Tid1 for an 84-mer ssDNA is about 230 ± 17 nM (Figure S5) which is about 4,600 fold lower than that of RPA at high-affinity mode (Kd ~0.05 nM) with a binding site size of ~30 nucleotides(51). Therefore, RPA is very likely to improve D-loop yields in the pre-mixed protocol by blocking inhibitory interactions of Rdh54/Tid1 with ssDNA, as for Hop2-Mnd1.

**Figure 5.**
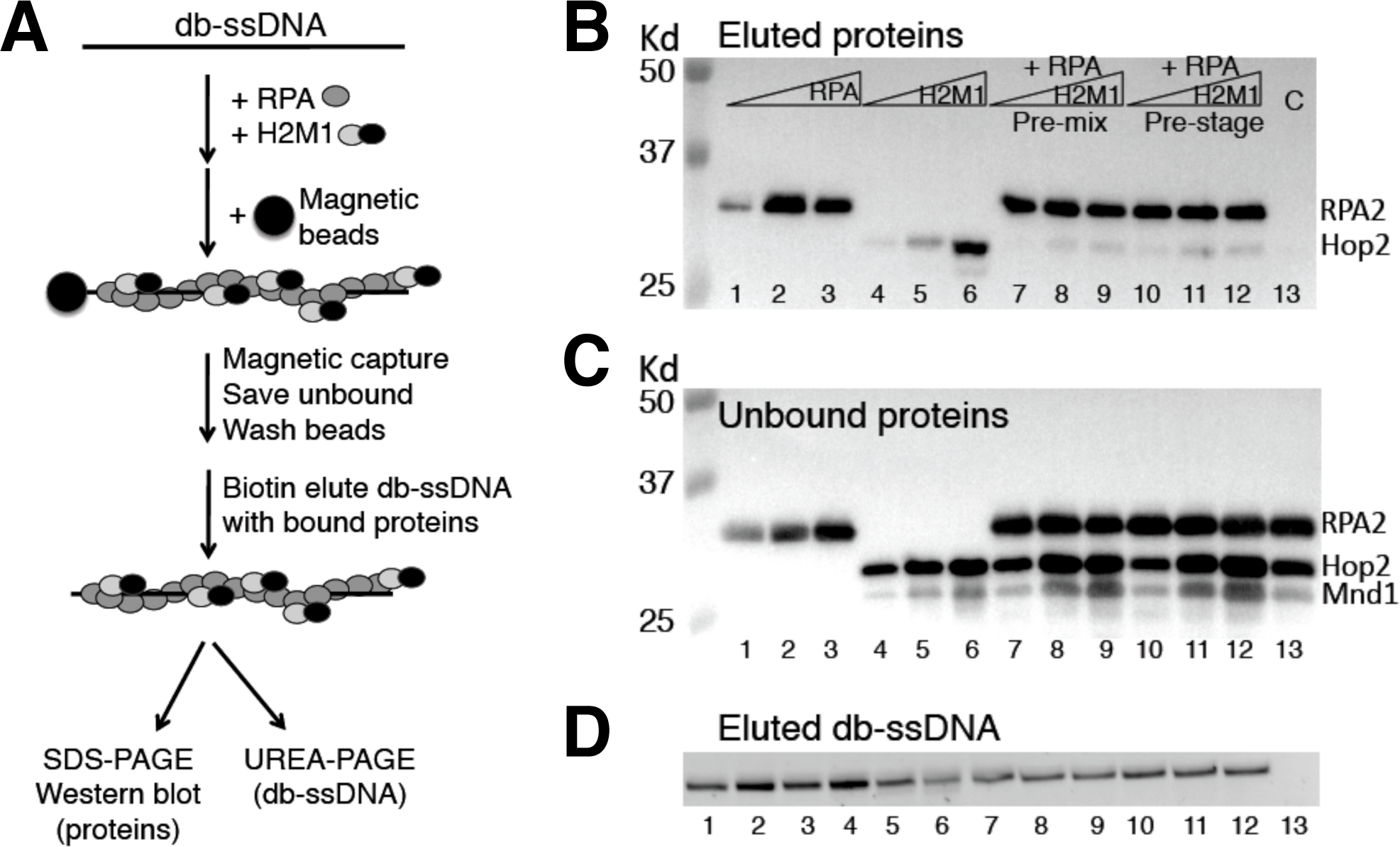
RPA outcompetes Hop2-Mnd1 for binding to ssDNA. (**A**) Scheme for bead “catch and release” of protein complexes bound to ssDNA. db-ssDNA stands for desthiobiotin modified ssDNA. (**B**) The captured protein complexes on ssDNA were analyzed by Western blotting. RPA were 0.125, 0.25, and 0.5 μM in lanes 1 - 3; 0.5 μM in lanes 7 -13. Hop2-Mnd1 were 0.2, 0.4, and 0.6 μM in lanes 4 - 6, lanes 7 -9, and lanes 10 -12. Lane 13 is a control that lacked db-ssDNA but contained 0.5 μM RPA and 0.4 μM Hop2-Mnd1. (**C**) The unbound protein complexes analyzed by Western blotting. (**D**) The eluted db-ssDNA-protein complexes were denatured and analyzed for DNA content on 8% urea-PAGE via staining with SYBR gold.

### Multiple protein-proteins interactions involving RPA, Dmc1, Rad51, and Mei5-Sae3

A “hand-off” model was proposed previously to account for RPA function; RPA binds other DNA processing proteins to order and guide the activities of the proteins that trade places on ssDNA (7). However, “hand-off’ mechanisms remain poorly understood. To gain a better understanding of how RPA exerts its functions during our *in vitro* reconstitution reactions, we analyzed the interactions of RPA with Dmc1 and its accessory proteins. Pairwise interactions with RPA were examined using Dmc1, Rad51-II3A, Rdh54/Tid1, Mei5-Sae3 and Hop2-Mnd1. The approach took advantage of the fact that all proteins used in the study, with the exception of RPA, carry (His_6_) tags allowing use of (His_6_)-Ni affinity pulldowns. Each of these proteins was incubated individually with RPA; then Ni resin was added to the mixture to capture protein or protein complexes through the (His_6_) tag. We found Dmc1 and Mei5-Sae3 each pulled down RPA (Figure 6A, lanes 2 and 3), but Rad51-II3A, Hop2-Mnd1, and Rdh54/Tid1 did not (Figure 6A, lanes 4 - 6). To verify these findings, we used a reciprocal method of *in vitro* co-immunoprecipitation by antibodies against RPA2 to pulldown proteins that interact with RPA. We confirmed the above physical interactions between RPA and Dmc1, and between RPA and Mei5-Sae3 (Fig. 6b, lanes 7, 8, 10). Although we did not detect interaction between RPA and Rad51 (Figure 6B, lane 9), Rad51 was pulled down with RPA if Mei5-Sae3 and/or Dmc1 were also present, with the highest level of Rad51 recovered when both were present (Figure 6B, lanes 12 - 14). To further verify these protein-protein interactions, we carried out co-immunoprecipitation using antibodies against Dmc1 (Figure 6C). Importantly, we detected direct interaction between Dmc1 and Rad51 (Figure 6C, lanes 6), and a previously described interaction between Dmc1 and Mei5-Sae3 (13, 15) (Figure 6C, lanes 8). Moreover, we showed Mei5-Sae3 enhanced the interaction between Dmc1 and Rad51 (Figure 6C, lanes 7). The results provide evidence for a network of interactions amongst these yeast proteins. RPA physically interacts with Dmc1 and Mei5-Sae3, and a complex of RPA-Dmc1-(Mei5-Sae3) contacts Rad51 through interactions involving Dmc1 and Mei5-Sae3.

**Figure 6.**
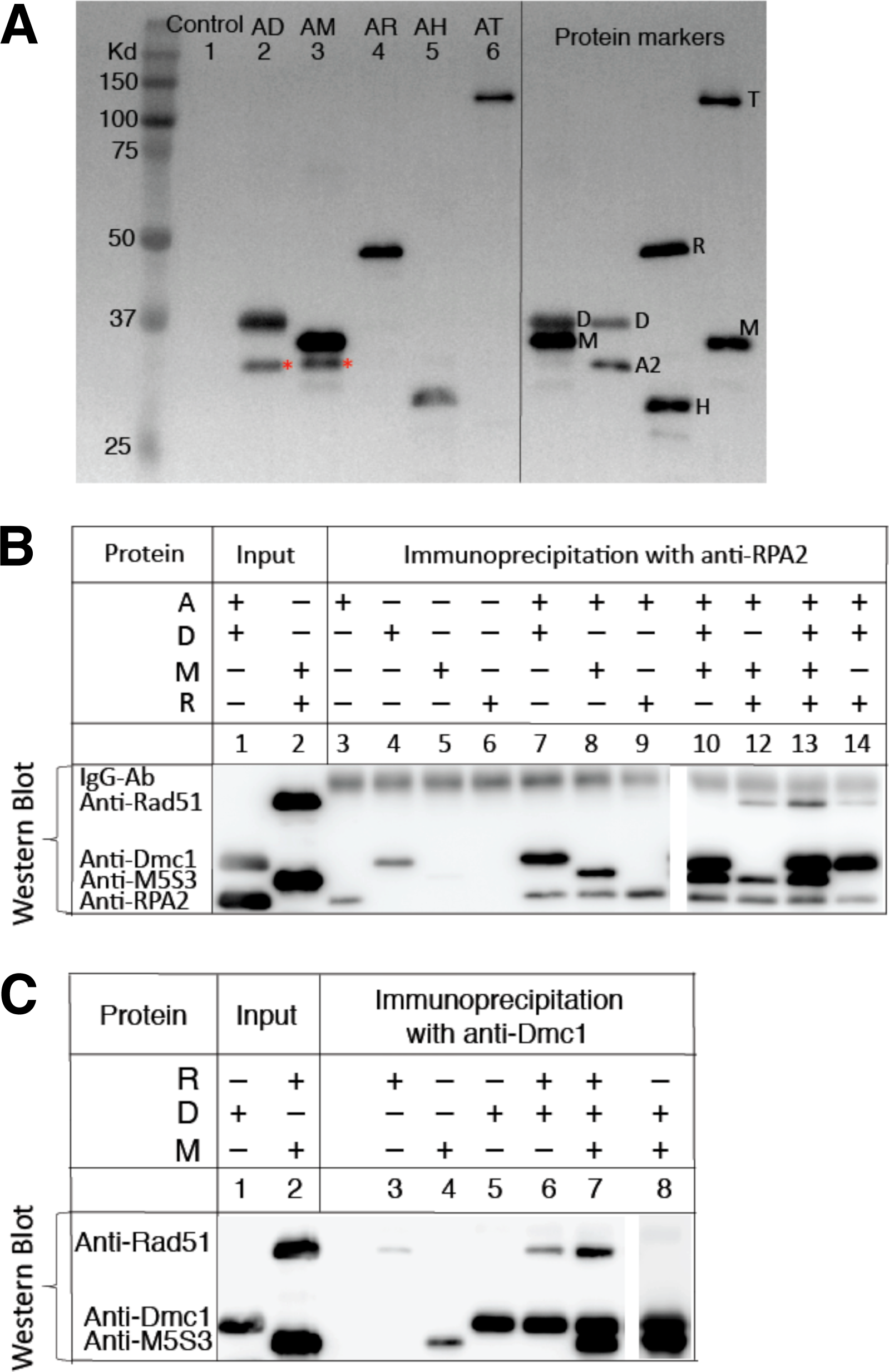
Protein-protein interactions. (**A**) Detection of direct protein-protein interactions by His6-affinity pulldowns: purified Dmc1 (D), Mei5-Sae3 (M), Rad51-II3A (R), Hop2-Mnd1 (H), and Rdh54/Tid1 (T) were (His)_6_-tagged. Each of these proteins was first incubated individually with untagged RPA, followed by the addition of Ni sepharose beads to capture the interacting protein complex through the (His)_6_-tag. The Ni captured complexes were eluted with a solution containing 1.7% SDS and 0.1 M DTT and separated on 12% SDS-PAGE, further analyzed by Western blotting using antibodies against all the proteins involved in the analysis for detection. An asterisk indicates the position of RPA2 (A2) pulled down by Dmc1 and Mei5-Sae3. (**B**) Detection of direct protein-protein interactions by *in vitro* co-immunoprecipitation of anti-RPA2 antibody with purified proteins as indicated. (**C**) Detection of direct protein-protein interactions by *in vitro* co-immunoprecipitation with anti-Dmc1 antibody. A mixture of antibodies against all proteins was used for protein detection.

### Physiological concentrations of meiotic recombination proteins

To compare our reconstituted conditions with relative protein levels *in vivo* we determined *in vivo* protein concentrations by quantitative Western blotting. Yeast whole cell extracts were prepared from synchronous meiotic cultures at various times following meiotic induction (Figure S6A and S6B). Peak protein concentrations were calculated, under the assumption that all proteins are localized to the nucleus (Table 1, Supplementary Figure S6C). The peak protein concentrations showed that RPA and Hop2 are the most abundant, and Rdh54/Tid1 the least. After normalizing the protein levels to that of Rdh54/Tid1, we found that the relative *in vivo* levels of Rdh54/Tid1, Rad51, Mei5/Sae3, and Dmc1 were within 3-fold of the relative levels of the same proteins under optimal D-loop conditions. The relative concentrations of RPA and Hop2-Mnd1 were much higher than those in the reconstituted system (see DISCUSSION).

## DISCUSSION

### Reconstitution of Dmc1-dependent D-loop formation

We have reconstituted meiotic D-loop formation using six purified *Saccharomyces cerevisiae* proteins including the strand exchange protein Dmc1 and its accessory proteins RPA, Rad51-II3A, Mei5-Sae3, Hop2-Mnd1, and Rdh54/Tid1. We found that, when added as a protein mixture to DNA substrates, each accessory protein contributes to the yield of D-loop products. The reconstitution conditions we identified support a much higher level of D-loop activity than previously reported, with up to 35% of duplex substrate converted to products. *In vivo* measurements of the steady state levels of these proteins indicate that, although the absolute concentrations at which the activities of the proteins are optimal in *our* biochemical experiments are much lower than their steady state concentrations *in vivo*, the relative optimum concentrations *in vitro* display significant similarity to the relative concentrations of the same proteins in wild type cells. This similarity suggests that the reconstitution conditions are biologically relevant. Two exceptions to the correspondence of relative protein levels *in vivo* and *in vitro* were Hop2-Mnd1 and RPA, which are present at much higher relative concentrations *in vivo.* We speculate that the effective concentration of Hop2-Mnd1 in the vicinity of a particular DNA recombination event is much lower than the overall nuclear concentration as a consequence of sequestration of the majority of protein at distant, non-recombining chromosomal sites. Our speculation is supported by previous cytological studies, as described above. The assumption that each protein species is fully localized to the nucleus is clearly problematic for RPA as a significant fraction of total cellular protein localizes to the cytoplasm (52, 53). If we assume that RPA is uniformly distributed in the cell, there is good agreement between the relative levels of RPA in the reconstituted system and in the nucleus. Further studies are required to determine the true nuclear concentration of RPA.

Our progress in biochemical reconstitution of meiotic recombination builds a strong foundation for future work. This work will be directed at improving the similarity between properties of the reconstituted system and those of Dmc1-mediated recombination *in vivo.* One significant way in which the current biochemical system differs from the *in vivo* situation is that mutant cells lacking either Hop2-Mnd1 or Mei5-Sae3 have almost no residual Dmc1-dependent D-loop activity, while omitting either protein from the reconstituted system causes only a modest reduction of Dmc1 activity. This suggests that *in vivo* conditions are more restrictive to D-loop formation than our *in vitro* conditions. An obvious possibility to account for the relative permissiveness of the *in vitro* system is that duplex DNA substrates are packaged in chromatin *in vivo*, but not *in vitro.* Therefore, future studies will employ chromatinized plasmids to determine if they impose more strict requirements for accessory proteins than those observed with naked DNA.

### RPA plays multiple roles in stimulating Dmc1

A particularly important observation that arose from our effort to optimize the efficiency of Dmc1-dependent D-loop formation is that RPA is critical for the efficiency of the process. Our experiments provide evidence for three distinct mechanisms that contribute to the RPA’s stimulatory activity. RPA stimulates D-loop formation by: (1) eliminating secondary structure (46, 47); (2) binding the displaced strand of the D-loop (44, 45); (3) a novel mechanism that prevents inhibitory interactions of Hop2-Mnd1 and Rdh54/Tid1, which occur during Dmc1 binding to ssDNA. At least one cause of accessory protein conflict is binding to ssDNA, which is prevented by RPA’s ability to outcompete the other proteins for binding to ssDNA.

### Protein-protein interactions associated with Dmc1 filament assembly

Protein-protein interactions have been found to induce conformational rearrangement of RPA’s modular domains in a manner that alters the mode of RPA-ssDNA binding resulting in hand-offs of ssDNA from RPA to other ssDNA binding proteins (7). As a first step towards examining this potential mechanism for RPA’s activity, we showed that RPA binds directly to Dmc1, demonstrating evolutionary conservation of an interaction previously reported for the corresponding mammalian orthologues (54).

Previous *in vivo* studies suggested that both Rad51 and Mei5-Sae3 act to recruit or stabilize Dmc1 at sites of recombination *in vivo* (13, 14) and that Rad51 and Mei5-Sae3 can cooperate to stimulate Dmc1 ‘s activity *in vitro* (11). However, the biochemical mechanism underlying the influence of Rad51 and Mei5-Sae3 on Dmc1’s activity has yet to be determined. Previous studies showed that Dmc1 binds Mei5-Sae3, Mei5-Sae3 binds Rad51, and RPA binds Mei5-Sae3 (13, 15). Here we report detection of protein-protein interaction between Rad51 and Dmc1. A conventional two-hybrid system failed to detect this interaction (our unpublished results), and a recent biochemical study showed that Rad51-Rad51 and Dmc1-Dmc1 homotypic interactions are strong enough to result in predominance of homofilaments when the two are allowed to bind DNA as a mixture (55). Cytological observations also argue for a predominance of homofilaments (12, 30, 56). Therefore, detection of direct Rad51-Dmc1 binding is significant because it provides a molecular explanation for the role of Rad51 in stimulating Dmc1 activity. We also show that Mei5-Sae3 can enhance interaction between Rad51 and Dmc1, consistent with our previous work that showed Rad51 and Mei5-Sae3 cooperate to stimulate Dmc1’s D-loop activity (11).

### Model

Our results may be incorporated into a model for the functions of Dmc1 accessory proteins that involves roles of RPA during both pre-synaptic filament formation and during strand exchange (Figure 7). The model incorporates many previous findings with those presented here. The steps in the model are as follows. (1) RPA binds ssDNA with high affinity, coating ssDNA. RPA removes secondary structures from ssDNA and blocks Hop2-Mnd1 and Rdh54/Tid1 from accessing ssDNA. (2) Protein-protein interactions involving Dmc1, Rad51, Mei5-Sae3 and RPA contribute to the assembly of functional Dmc1 filaments, with interaction of Rad51 and Dmc1 stabilized by protein-protein interactions with Mei5-Sae3 (57). (3) Hop2-Mnd1 stabilizes non-homology dependent interactions between Dmc1-ssDNA filaments and dsDNA (26, 58, 59). (4) Rdh54/Tid1 stabilizes nascent loops by virtue of its ability to simultaneously bind both a ssDNA and a dsDNA molecule and/or its translocase activity (23)(Ref. 23 and references therein). (5) as D-loops form, RPA binds the displaced strand, blocking D-loop dissociation and extending D-loop length.

**Figure 7.**
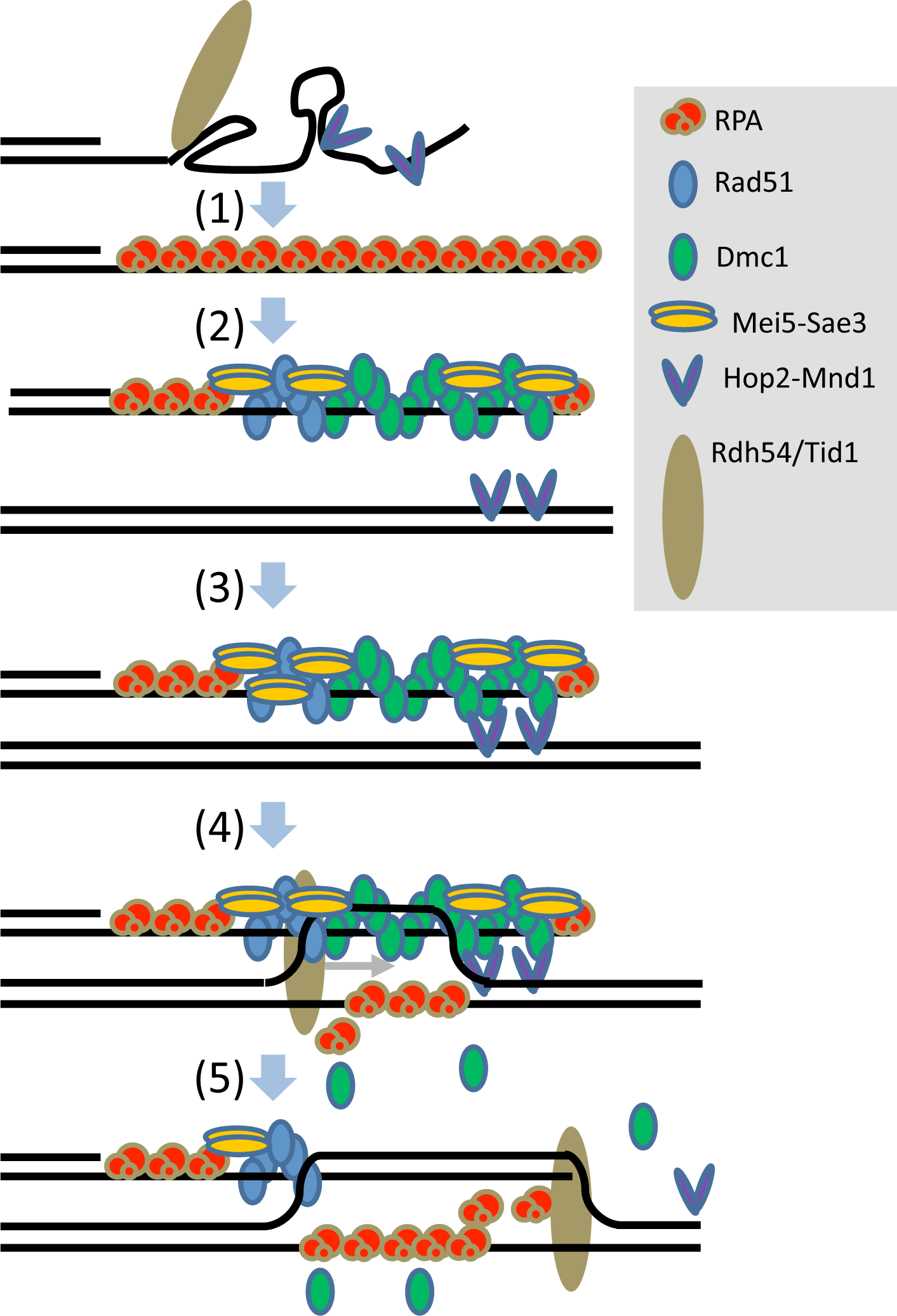
Model (1) RPA binds ssDNA thereby removing secondary structure and preventing binding of Hop2-Mnd1 and Rdh54/Tid1. (2) RPA carries out hand-off reactions allowing Rad51+Mei5-Sae3 to assist Dmc1 filament formation. The protein configuration shown is speculative but depicts potentially relevant protein-protein interactions between RPA and Mei5-Sae3, RPA and Dmc1, Rad51 and Mei5-Sae3, Dmc1 and Mei5-Sae3, and, particularly importantly, between Dmc1 and Rad51. (3) dsDNA bound Hop2-Mnd1 captures filaments via homology-independent interactions with Dmc1. (4) Homology recognition occurs to form a homology-dependent nascent D-loop which is stabilized by RPA binding to the displaced ssDNA strand. (5) Rdh54/Tid1 binds the D-loop, via contacts with both ssDNA and dsDNA and then translocates across the D-loop stabilizing it by displacing Dmc1 and elongating the D-loop. RPA can elongate the D-loop, even in the absence of Rdh54/Tid1, but does not afford the stability associated with Dmc1 displacement and/or simultaneous binding by Rdh54/Tid1 to both ssDNA and dsDNA.

### Conclusion

We report major progress in reconstitution of meiotic recombination reactions by identifying conditions under which a set of five accessory proteins cooperate to enhance the efficiency of Dmc1-mediated D-loop formation. Our results demonstrate the critical importance of RPA to the system, with evidence for three mechanisms of RPA-mediated stimulation of Dmc1-depednent D-loop reactions. The mechanisms of RPA stimulation include one that is particularly important and previously undescribed; RPA resolves inhibitory and conflicting activities of Hop2-Mnd1 and Rdh54/Tid1. We also detect novel protein-protein interactions including direct interaction between Rad51 and Dmc1, a result that further supports the view that meiotic recombination events require cooperation of Rad51 and Dmc1, with Dmc1 serving as enzyme, and Rad51 as accessory protein.

## SUPPLEMENTARY DATA

Supplementary figures and legends are available at NAR Online.

## ACKNOWLEDGMENTS

We thank Diedre Reitz and Brian Budke for comments on the manuscript, and Joseph Piccirilli for advice on analysis of kinetic data. We thank A. Shinohara (Osaka University, Japan) for providing antibodies against RPA2.

## SUPPLEMENTARY FIGURES AND LEGENDS

**Figure S1.**
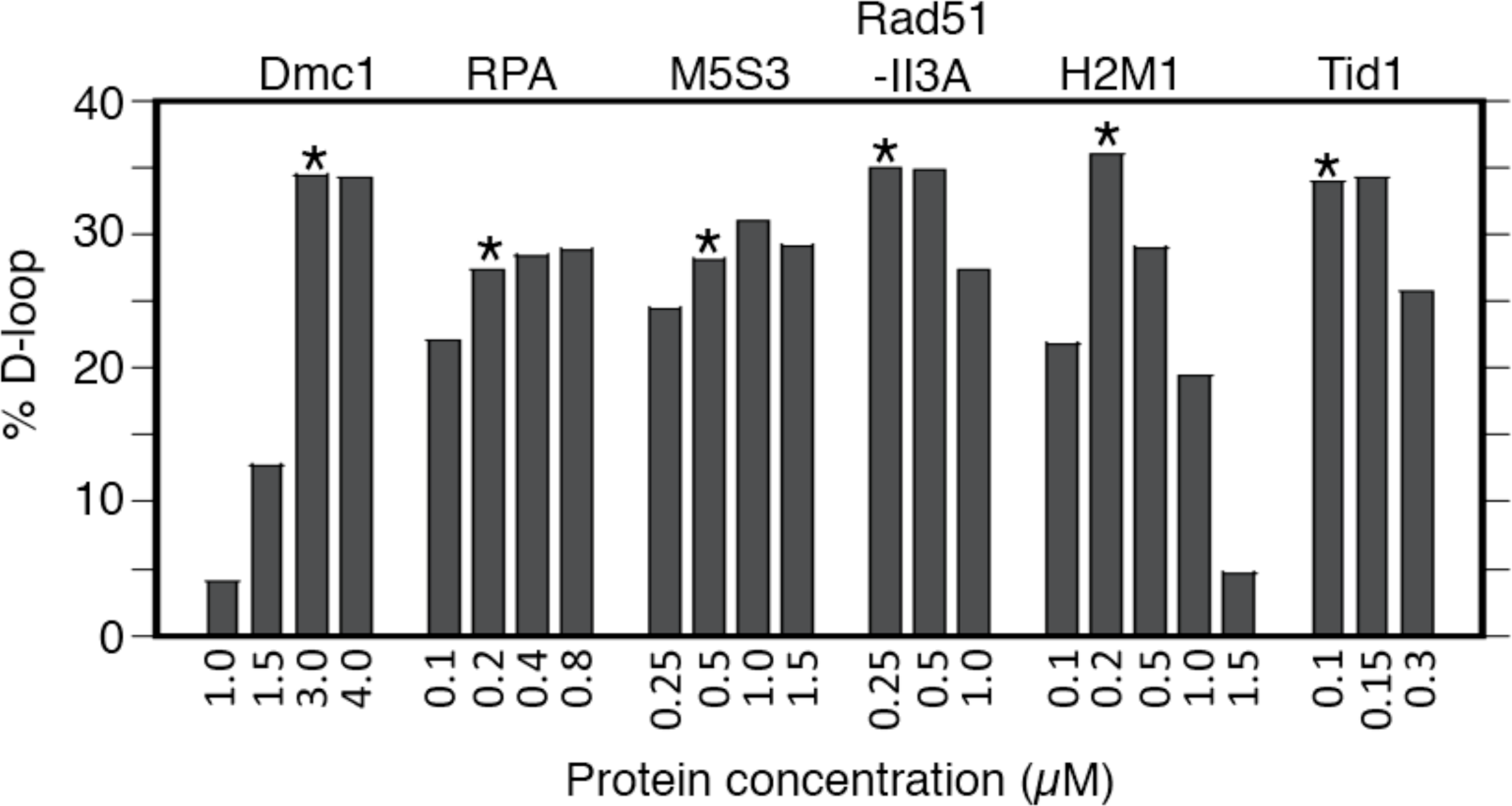
Determination of optimal protein concentration in the reconstitution reaction by titration. Titration was done in the D-loop assay using the 563 nt ssDNA (4 nM or 2.2 μΜ nt) and all proteins. Conditions were as described in Figure 1. In each set of titration reactions, only the concentration of one protein was varied as indicated in the figure, the concentrations of the remaining proteins were kept constant. The concentration indicated by an asterisk was used in all experiments presented in this paper, unless otherwise indicated.

**Figure S2.**
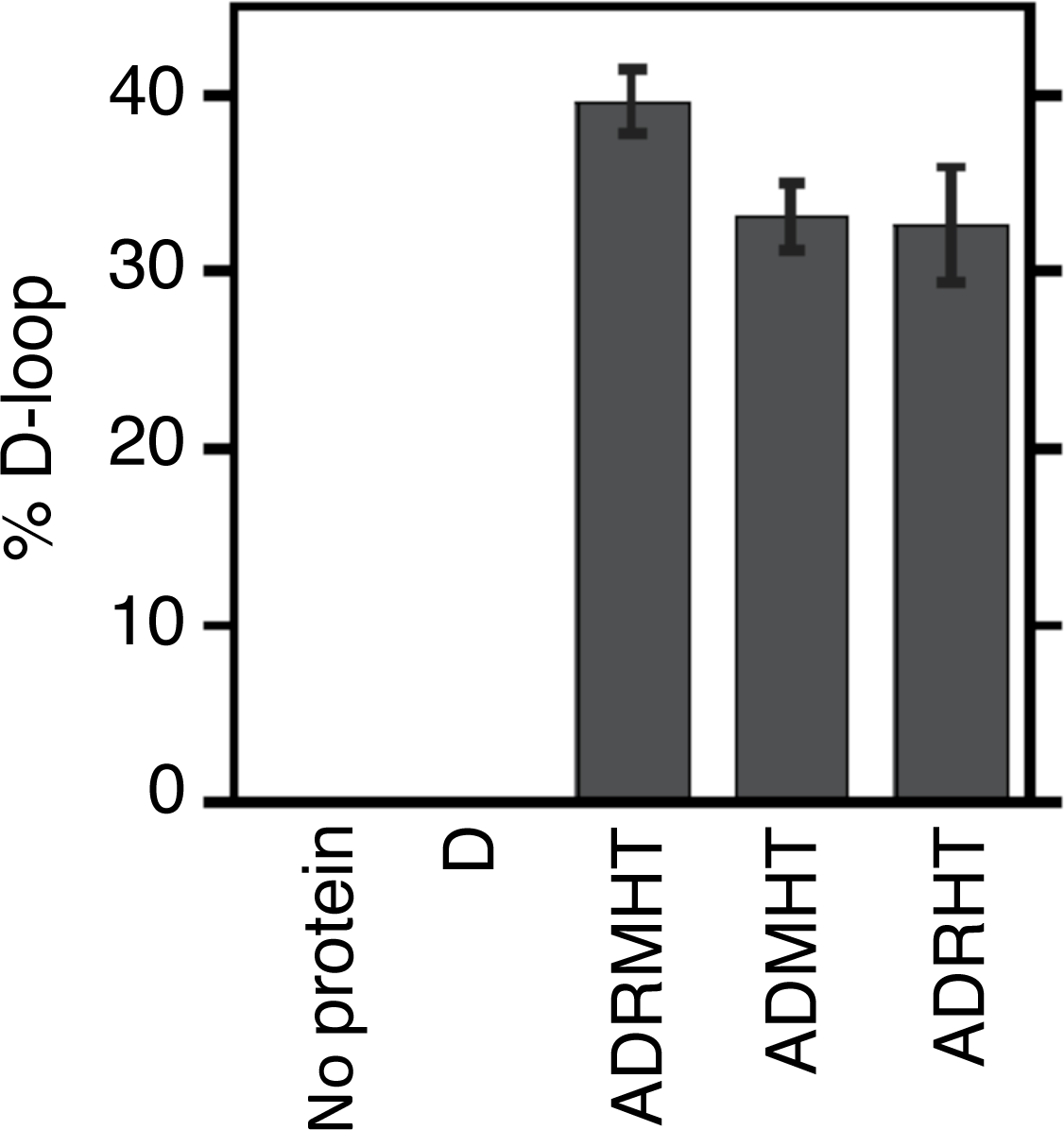
Mei5-Sae3 and Rad51 contribute to stimulating Dmc1’s D-loop activity in reconstitution. This experiment was necessary because we added or omitted a combination of Mei5-Sae3 and Rad51 in the experiments shown in Figure 1 for feasibility. Conditions were as in Figure 1.

**Figure S3.**
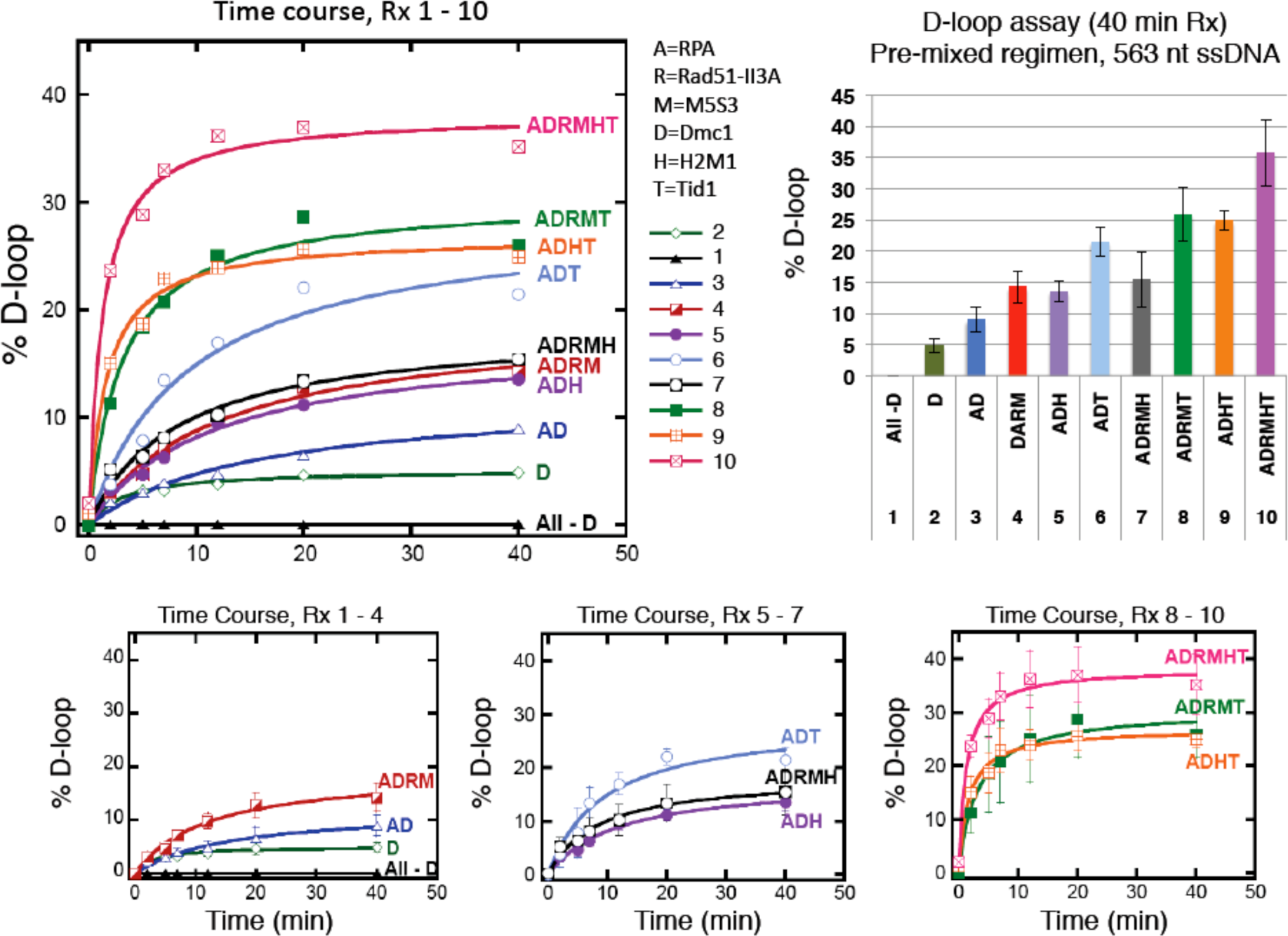
Time course of D-loop formation in reconstitution reactions. Dmc1’s D-loop formation as a function of time was determined in D-loop assay using the 563 nt ssDNA (4 nM or 2.2 μM nt) and various combinations of accessory proteins as indicated in the pre-mixed regimen. The D-loops levels at 0, 2, 5, 7, 12,and 40 min were plotted (n = 3, ± s.e.m). The time course of reactions numbered 1 - 10 were first plotted without error bars to avoid overcrowding of data points. Then the time courses were replotted in separate panels to show error bars. The D-loop formed at the end-point of the time courses at 40 min were shown as bar plot. All curves and bar plots were color coded to represent proteins in various combinations as indicated.

**Figure S4.**
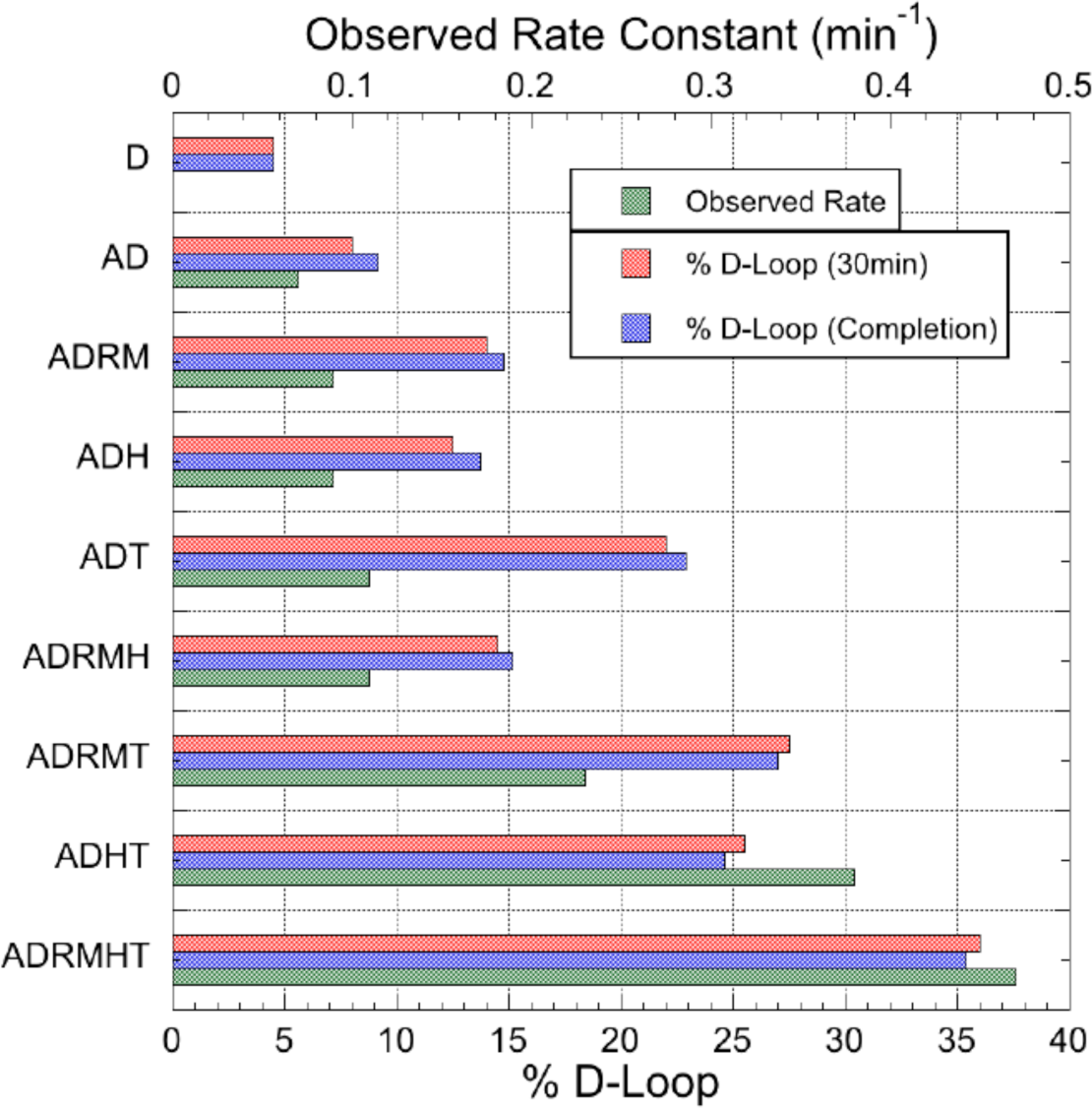
The apparent rate constant *(k*_*obs*_) of synaptic complex formation determined from reconstitution time course reactions in Supplementary Figure S3. Abbreviations for proteins: A = RPA; D = Dmc1; R = Rad51-II3A; M = Mei5-Sae3; H = Hop2-Mnd1; T = Rdh54/Tid1. (1) Data taken from time-courses in Supplemental Figure S3. (2) Endpoint (% D-Loop at t_∞_) and *k*_obs_ were determined by fitting time-course data to a simple kinetic model: y = A (1 - *e* ^−*kt*^). Where A is the endpoint and *k* is an apparent first-order rate constant. Data were fitted in Kaleidagraph™ (3) Reaction with only Dmc1 could not be fitted to the kinetic model because it had reacted to near completion by the first time-point.

**Figure S5.**
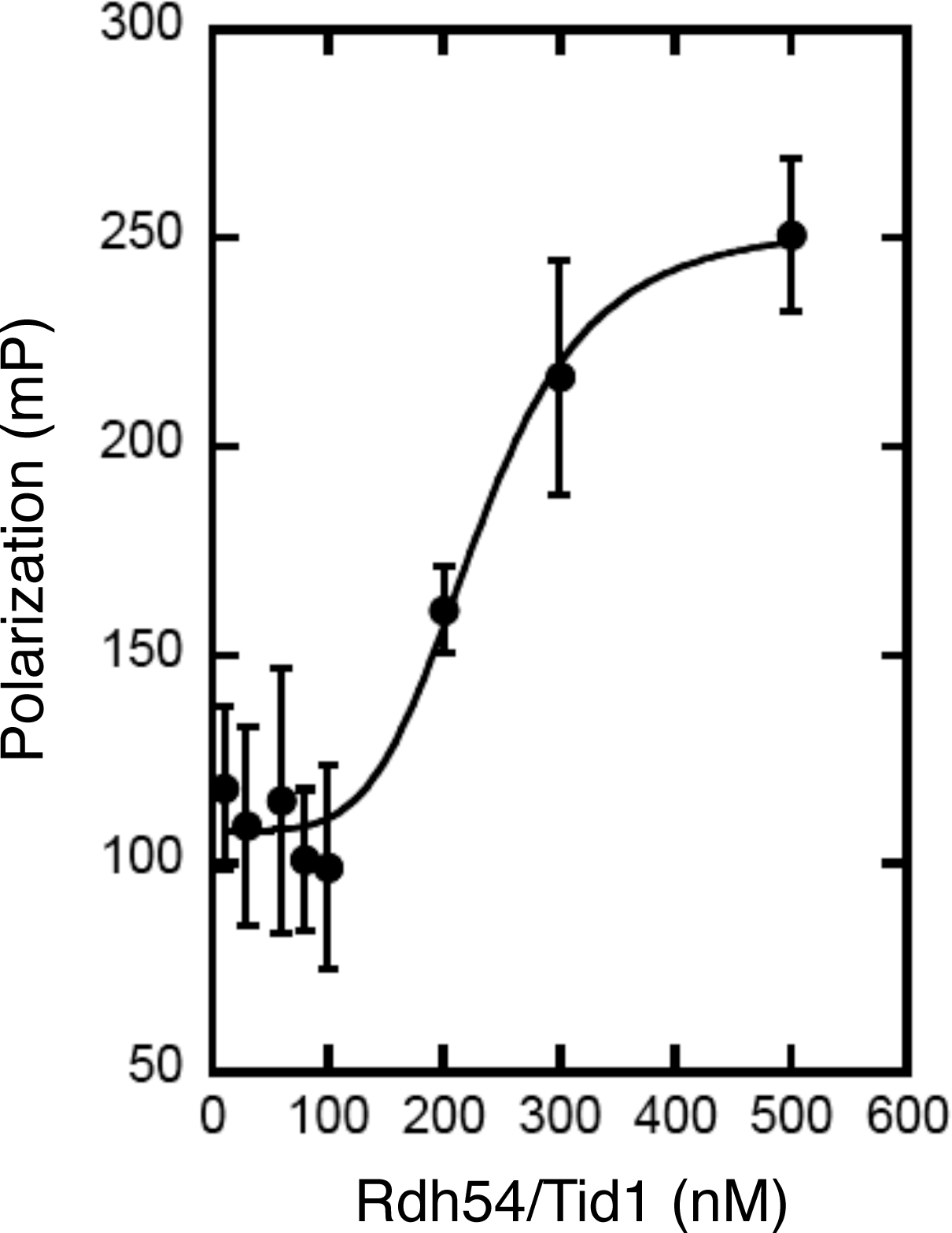
Rdh54/Tid1 binds ssDNA as revealed by fluorescence polarization. Protein binding was to Alexa Fluor-488 labeled 84mer ssDNA using the polarization assay essentially as described previously (60). mP indicates millipolarization unit. The apparent affinity of Rdh54/Tid1 for ssDNA is 230 ± 17 nM (n = 3, ± s.e.m).

**Figure S6.**
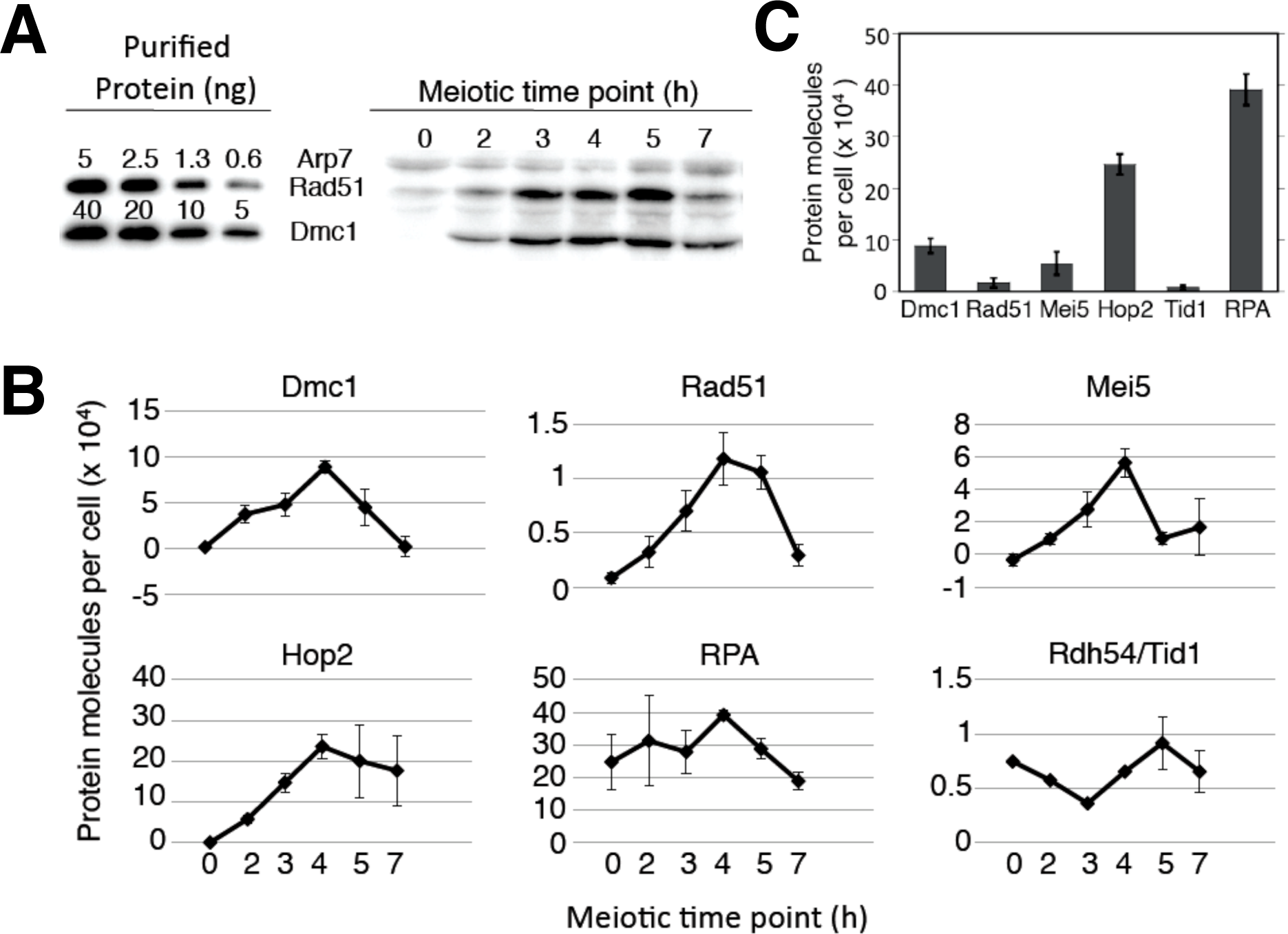
Physiological concentrations of proteins in budding yeast during meiosis. (**A**) An example Western blot. The protein concentrations of Dmc1, Rad51, Mei5, Hop2, Rdh54/Tid1, and RPA in meiotic budding yeast were determined by the Western blotting method. Total cellular proteins extracted from meiotic cells at the times indicated were analyzed next to a dilution series of purified Rad51 and Dmc1. Both purified proteins contained (His_6_)-tag and thus migrated slightly slower in SDS-PAGE. Detection was with antibodies against Rad51, Dmc1, and Arp7. Arp7 is known to have stable expression during meiosis and was used as an internal standard for band normalization. (**B**) The expression profiles of 6 proteins. The mean values of each time point from 3 independent time courses were plotted and error bars show ± s.e.m. (**C**) A bar plot comparing the number of protein molecules per yeast meiotic cell of 6 proteins at their respective peak expression levels from **B**.

